# Reward Circuit Adaptations After Chronic Antipsychotic Treatment Confer Addiction Vulnerability

**DOI:** 10.64898/2026.04.24.720671

**Authors:** Anna Kruyer, Ariana Angelis, Connor Waselenko, Aaron Tan, Shawn Xiong, Davide Amato

**Author notes:** Authors contributed equally (Anna Kruyer, Ariana Angelis). **Location of Work:** Work presented herein was completed at the University of Cincinnati, Cincinnati, OH and the Medical University of South Carolina, Charleston, SC. **Correspondence:** Address correspondence to Davide Amato, P.D., Ph.D., 231 Albert Sabin Way, MSB 3005, Cincinnati, OH 45267 or Anna Kruyer, Ph.D., 231 Albert Sabin Way, MSB 3005, Cincinnati, OH 45267.

## Abstract

**Objective:** Antipsychotics are widely used chronically to treat several psychiatric conditions and off-label indications, but their clinical utility is often accompanied by motor and/or metabolic side effects. Antipsychotics can also alter the brain’s reward circuitry, and we recently showed that upon abrupt discontinuation, chronic treatment with the first-generation antipsychotic haloperidol increases cocaine relapse in a rodent model of i.v. cocaine self-administration. Here, we examined whether chronic treatment with second-generation antipsychotics was also associated with increased addiction vulnerability in rodents and humans, and sought to further characterize the cellular substrate linking antipsychotic exposure to substance use disorder (SUD).

**Methods:** Adult rats received subcutaneous infusion of risperidone, olanzapine, or vehicle for 14 days. Rats were then trained to self-administer i.v. cocaine or oral sucrose pellets for 10 days and active lever presses were paired with compound cue exposure. After a period of extinction training, reward seeking was reinstated by restoration of cues to active lever pressing for 20 minutes. To enable quantification of dendritic spine morphology and manipulation of MSNs during relapse, viral approaches were used to label D1- or D2-MSNs in the nucleus accumbens core (NAcore), a region that encodes reward-related responses and regulates symptoms associated with addiction and other psychiatric disorders, including schizophrenia. To assess the clinical relationship between chronic antipsychotic use and SUD, we conducted a systematic review and meta-analysis of studies identified through PubMed, CENTRAL, Scopus, and Embase that examined the effects of antipsychotic treatment on addictive substance use or craving in patients with SUD.

**Results:** In rats, chronic risperidone or olanzapine treatment delayed extinction learning and increased cue-induced cocaine seeking, but did not alter seeking of the natural reward sucrose or its extinction. Cued relapse increased spine head diameter in NAcore D2-MSNs, but not D1-MSNs after antipsychotic treatment relative to vehicle-treated controls. Chemogenetic inhibition of D2-MSNs reversed addiction vulnerability conferred by antipsychotic treatment. In patients whose SUDs were treated with antipsychotics, duration of treatment (>4 months) strongly predicted addictive substance use or craving independent of antipsychotic type (first- vs. second-generation) or misused substance (psychostimulant, opioid, alcohol, or cannabis).

**Conclusions:** Chronic antipsychotic treatment, regardless of drug class, was associated with increased vulnerability to addictive substance use and relapse-like behavior in a rat model, and was linked to potentiation of D2-MSNs in the ventral striatum. In humans, systematic review and meta-analytic findings supported this pattern in patients maintained on antipsychotic treatment.

## Introduction

Substance use disorders (SUDs) are substantially more common among individuals with mental illness, representing one of the most consistent findings in psychiatric epidemiology (1). Roughly 50% of individuals diagnosed with a SUD meet criteria for at least one additional psychiatric disorder, far exceeding rates observed in the general population (2, 3). Similarly, up to 50% of individuals with a serious mental illness, including schizophrenia or bipolar disorder, develop a SUD over the course of a lifetime (4). High rates of co-occurrence have been documented across a wide range of psychiatric diagnoses, including major depression and bipolar disorder (5–7), ADHD (8, 9), schizophrenia and related psychotic disorders (2, 3), borderline personality disorder (10), and antisocial personality disorder (11). In individuals with psychotic disorders, comorbid SUDs can destabilize illness course and are linked to higher rates of hospitalization, homelessness, aggression, violence, incarceration, and suicidality (12–18). They can also worsen cognitive and negative symptoms, impair treatment response, reduce medication adherence, and increase the likelihood of relapse (14, 19–22). Understanding the factors that underlie this co-occurrence is critical for defining causal relationships between the two disorders and for developing strategies to prevent substance dependence in individuals with schizophrenia (23).

The most widely accepted explanations for the high comorbidity between psychotic disorders and SUDs emphasize shared vulnerability factors, including genetics, early-life adversity, and overlapping neurobiological mechanisms that increase risk for both SUDs and other psychiatric illnesses (13, 24). Additional hypotheses include the possibility that patients with psychiatric diagnoses self-medicate certain symptoms with addictive substances or are more likely to engage in high-risk behaviors, including addictive substance use (25–28). Instead, preclinical data suggest that, under certain conditions, chronic exposure to medications used to treat mental illness, including first- and early second-generation antipsychotics, may itself represent a risk factor for increased vulnerability to addictive substance use and relapse (29, 30). In support of this alternative explanation, several studies indicate that antipsychotic treatment, particularly with first-generation agents, alters reward processing and can blunt normal reward responsiveness (31–34). These changes may promote the use of addictive substances, especially psychostimulants, as a means of compensating for reduced striatal dopamine (34–37), which may underlie elevated excitatory signaling in the brain’s nucleus accumbens core (NAcore) that drives drug seeking during periods of abstinence (30).

Despite these strong links, there is no direct evidence supporting a definitive causal relationship between antipsychotic use in patients and SUD severity, nor is there clear preclinical or clinical evidence that links second-generation antipsychotics with SUD vulnerability. Instead, the clinical literature on this topic remains difficult to interpret, with inconsistent findings due to heterogeneous study designs and patient populations. Importantly, while patients with SUDs are often excluded from antipsychotic efficacy trials for methodological reasons (38, 39), withholding antipsychotic treatment from patients with psychotic disorders is unethical, precluding direct assessment of the causal effects of antipsychotics on SUDs-related outcomes relative to placebo control treatment. Thus, given the prevalence of addictive substance use in schizophrenia, this remains an important yet inherently difficult area to study. To begin addressing these critical questions, we conducted a comprehensive set of studies in an animal model of cocaine addiction to identify potential underlying mechanisms and, in parallel, performed a systematic review and meta-analysis to examine whether antipsychotic treatment was associated with increased addiction liability in humans as suggested by preclinical studies.

## Methods

### Antipsychotic Treatment

Experimental procedures involving animals were conducted in accordance with guidelines established by the Institutional Animal Care and Use Committees at the University of Cincinnati and the Medical University of South Carolina. Male and female D1- or D2-Cre rats or non-transgenic littermates were bred in house and maintained on a reverse 12:12 light:dark cycle and were fed *ad libitum* throughout experimentation. Rats (175-200g) were implanted with subcutaneous osmotic infusion pumps (Alzet, LLC) delivering olanzapine (2.5 mg/kg/d), risperidone (4 mg/kg/d) or vehicle for 14 days. After 14 days, pumps were removed under isoflurane anesthesia. Antipsychotic washout occurred for 7 days prior to initiation of cocaine self-administration.

### Operant Self-Administration, Extinction and Reinstatement

After osmotic pump removal, rats were implanted with intrajugular catheters and after recovery from surgery, rats were trained to self-administer i.v. cocaine (0.4 mg/kg/infusion) during 2-hour daily sessions for 10 days. Animals trained to self-administer sucrose did not undergo catheter implantation and received oral sucrose pellets (45 mg, Bio-Serv) instead of i.v. cocaine infusions. Active lever presses for cocaine or sucrose were paired with reward delivery and light and tone cues for 5 seconds. After self-administration, animals underwent 12 days of extinction training (2 hours/day) where lever presses had no consequence. Lever pressing was reinstated by exposure to cues previously paired with reward delivery for 20 minutes 24 hours after the last extinction session. Reinstatement sessions were initiated by a single exposure to experimenter-delivered light and tone cues. During reinstatement sessions, active lever presses resulted in cue exposure, but no reward delivery.

### Dendritic Spine Morphology

During catheter implantation, D1- and D2-Cre rats received infusions of AAV2-CAG-Flex-Ruby2sm-Flag (1 μL/hemisphere, 0.15 μL/min, 5 min diffusion; Addgene, Inc.) in the NAcore (+1.5 mm AP, ±1.7 mm ML, -7.0 mm DV), a brain structure that processes reward and plays a key role in addiction and other psychiatric disorders, including schizophrenia (40). Virus incubation occurred over the course of operant training. After 20 minutes of cued reinstatement 24 hours after the last extinction session, or after 20 minutes of extinction training 24 hours after the last self-administration sesssion, animals were anesthetized with intrahepatic delivery of ketamine (5 mg) and perfused transcardially with 4% paraformaldehyde. Coronal slices (50 μm) containing the NAcore were collected using a vibratome (Leica Biosystems) and stored at 4C until use. Tissue slices were permeabilized in phosphate-buffered saline (PBS) with 2% Triton X-100 for 1-hour at room temperature and nonspecific epitope binding was blocked by incubation in PBS with 0.2% Triton X-100 (PBST) and 2% normal goat serum for 1 hour at room temperature before overnight incubation in primary antibody (mouse anti-FLAG, F1804, Sigma-Aldrich) at 1:1000 in block at 4C. After washing in PBST, tissue was incubated in fluorescent secondary antibody (Thermo Fisher Scientific) at room temperature for 2 hours before washing and mounting. Images were collected using a Leica Stellaris 5 using a 63x objective lens and 3.5x digital zoom with automated, adaptive post-capture processing (Lightning, Leica Microsystems). Images were collected using a 1024 x 256 frame size, 4-frame averaging, and a 0.21-μm step size. Dendritic segments were imaged 75-200 μm from the cell soma after branching. Following acquisition, z-series were exported to Imaris 9.8 (Oxford Instruments) for quantitation. Imaging and analysis were performed by an experimenter blind to animal treatment.

### D2-MSN Inhibition

During catheter implantation, D2-Cre rats received infusions of AAV2-hSyn-DIO-hM4D(Gi)-mCherry (1 μL/hemisphere, 0.15 μL/min, 5 min diffusion; Addgene, Inc.) in the NAcore using coordinates listed above and virus incubation occurred over the course of operant training. Control animals received intracranial saline infusion in lieu of virus delivery. Animals received CNO (3 mg/kg, i.p.; Tocris Bioscience) 15 minutes prior to cued reinstatement of seeking and active lever pressing was monitored for 20 minutes.

### Systematic Review and Meta-Analysis

Papers related to the impact of antipsychotic treatment on addictive substance use and craving were collected using the following search terms:

abilify[Title/Abstract] OR aceperone[Title/Abstract] OR aceprometazine[Title/Abstract] OR acetophenazine[Title/Abstract] OR amisulpride[Title/Abstract] OR antipsychotic*[Title/Abstract] OR aripiprazole[Title/Abstract] OR asenapine[Title/Abstract] OR benperidol[Title/Abstract] OR brexpiprazole[Title/Abstract] OR bromperidol[Title/Abstract] OR butaperazine[Title/Abstract] OR butyrophenone[Title/Abstract] OR carpipramine[Title/Abstract] OR chlorpromazine[Title/Abstract] OR chlorprothixene[Title/Abstract] OR clocapramine[Title/Abstract] OR clofluperol[Title/Abstract] OR clopenthixol[Title/Abstract] OR clotiapine[Title/Abstract] OR clozapine[Title/Abstract] OR compro[Title/Abstract] OR cyamemazine[Title/Abstract] OR “D2 antagonist”[Title/Abstract] OR “D-2 receptor”[Title/Abstract] OR “D2 receptor antagonist”[Title/Abstract] OR “D2/D3 antagonist”[Title/Abstract] OR “D2/D3 receptor antagonist”[Title/Abstract] OR “D2r antagonist”[Title/Abstract] OR “D3 antagonist”[Title/Abstract] OR “D-3 receptor”[Title/Abstract] OR “D3 receptor antagonist”[Title/Abstract] OR “D3r antagonist”[Title/Abstract] OR dixyrazine[Title/Abstract] OR droperidol[Title/Abstract] OR esucos[Title/Abstract] OR fluanisone[Title/Abstract] OR Fluanxol[Title/Abstract] OR flupenthixol[Title/Abstract] OR flupentixol[Title/Abstract] OR fluphenazine[Title/Abstract] OR fluspirilene[Title/Abstract] OR flutroline[Title/Abstract] OR haldol[Title/Abstract] OR haloperidol[Title/Abstract] OR iloperidone[Title/Abstract] OR invega[Title/Abstract] OR Largactil[Title/Abstract] OR lenperone[Title/Abstract] OR levomepromazine[Title/Abstract] OR loxapine[Title/Abstract] OR lurasidone[Title/Abstract] OR “major tranquiliz*”[Title/Abstract] OR melperone[Title/Abstract] OR mesoridazine[Title/Abstract] OR methopromazine[Title/Abstract] OR methotrimeprazine[Title/Abstract] OR metitepine[Title/Abstract] OR molindone[Title/Abstract] OR moperone[Title/Abstract] OR neuroleptic*[Title/Abstract] OR neurolepticum[Title/Abstract] OR Nozinan[Title/Abstract] OR olanzapine[Title/Abstract] OR orap[Title/Abstract] OR oxypertine[Title/Abstract] OR oxyprothepine[Title/Abstract] OR paliperidone[Title/Abstract] OR penfluridol[Title/Abstract] OR perazine[Title/Abstract] OR periciazine[Title/Abstract] OR pericyazine[Title/Abstract] OR perphenazine[Title/Abstract] OR “phenothiazine tranquiliz*”[Title/Abstract] OR piflutixol[Title/Abstract] OR pimavanserin[Title/Abstract] OR pimozide[Title/Abstract] OR pipamperone[Title/Abstract] OR piperacetazine[Title/Abstract] OR pipotiazine[Title/Abstract] OR prochlorperazine[Title/Abstract] OR promazine[Title/Abstract] OR propericiazine[Title/Abstract] OR propiomazine[Title/Abstract] OR prothipendyl[Title/Abstract] OR quetiapine[Title/Abstract] OR raclopride[Title/Abstract] OR remoxipride[Title/Abstract] OR reserpine[Title/Abstract] OR Risperdal[Title/Abstract] OR risperidone[Title/Abstract] OR Seroquel[Title/Abstract] OR sertindole[Title/Abstract] OR setoperone[Title/Abstract] OR spiperone[Title/Abstract] OR Stelazine[Title/Abstract] OR Stemetil[Title/Abstract] OR stepholidine[Title/Abstract] OR sulforidazine[Title/Abstract] OR sulpiride[Title/Abstract] OR taxilan[Title/Abstract] OR tetrabenazine[Title/Abstract] OR thiopropazate[Title/Abstract] OR thioproperazine[Title/Abstract] OR thioridazine[Title/Abstract] OR thiothixene[Title/Abstract] OR tiapride[Title/Abstract] OR tiotixene[Title/Abstract] OR trifluoperazine[Title/Abstract] OR trifluperidol[Title/Abstract] OR triflupromazine[Title/Abstract] OR ziprasidone[Title/Abstract] OR zotepine[Title/Abstract] OR zuclopenthixol[Title/Abstract] OR Zyprexa[Title/Abstract]

AND

((”narcotic*”[Title/Abstract] OR “opioid”[Title/Abstract] OR “heroin”[Title/Abstract] OR “morphine”[Title/Abstract] OR “opiate”[Title/Abstract] OR “opium”[Title/Abstract] OR “alcohol”[Title/Abstract] OR “tobacco”[Title/Abstract] OR “drug”[Title/Abstract] OR “cocaine”[Title/Abstract] OR “marijuana”[Title/Abstract] OR “amphetamine”[Title/Abstract] OR “substance”[Title/Abstract] OR “chemical”[Title/Abstract]) AND (”use”[Title/Abstract] OR “user*”[Title/Abstract] OR “abuse”[Title/Abstract] OR “dependence”[Title/Abstract] OR “misuse*”[Title/Abstract] OR “addict*”[Title/Abstract] OR “overdose”[Title/Abstract] OR “overuse*”[Title/Abstract])) OR alcoholi*[Title/Abstract]

OR

”substance disorder”[Title/Abstract] OR “substance disorders”[Title/Abstract] OR “alcohol disorders”[Title/Abstract] OR “alcohol disorder”[Title/Abstract].

We searched CENTRAL, Embase, Scopus and PubMed for studies until December 15, 2023. Title and abstract screening and full text screening were carried out independently by two researchers. Data extraction was carried out independently by two researchers with conflicts resolved by a third. Studies reported in English examining the effects of antipsychotic treatment on addictive substance craving or use were analyzed for qualitative and quantitative data. Extracted data were synthesized categorically, according to statistically significant results reported in each study, and quantitatively.

### Receptor Affinity

Receptor affinity of first- (haloperidol) and second-generation antipsychotics (olanzapine, risperidone) were calculated based on data collected and catalogued by the Psychoactive Drug Screening Program (National Institute of Mental Health). Multiplicate reports for each receptor or transporter subtype were averaged across studies with values and references reported in **Supplementary Data File 2**.

### Data Analysis

All experiments included groups with 4 or more animals each to minimize variability due to animal behavior. Biological replicates contained animals from all groups, including vehicle controls to minimize the impact of experimental variability on independent group values. No outliers were removed from any dataset. Data were analyzed using GraphPad Prism 10. All behavioral data were analyzed using one-tailed Student’s *t*-test or one-or two-way analysis of variance (ANOVA) with Bonferroni correction applied for multiple comparisons. Spine head diameter and human data were analyzed using Chi^2^.

## Results

### Chronic Treatment with Second-Generation Antipsychotics Increased Measures of Context- and Cue-Induced Cocaine Seeking in a Rodent Model of SUD

Adult Long Evans rats were treated with olanzapine (2.5 mg/kg/d), risperidone (4 mg/kg/d), or vehicle delivered subcutaneously via osmostic infusion for 14 days (**Fig. 1A**). These doses fall within the clinically-relevant range after appropriate adjustment for interspecies differences in body weight and drug metabolism. After a period of abstinence, rats were trained to self-administer i.v. cocaine (**Fig. 1A-B**) for 10 consecutive days and cocaine infusions were paired with light and tone cues. Rats then underwent 12 days of extinction training, where cocaine and cues were withheld in response to active lever pressing. While history of antipsychotic intake did not impact active lever pressing or cocaine intake during self-administration, olanzapine- and risperidone-treated animals pressed the active lever significantly more during extinction training compared to vehicle-treated control animals, despite absence of reward delivery (**Fig. 1C**), a finding that was observed in both male and female rats (**Fig. S1A-B**). During a cue-induced reinstatement session, a classic model of addictive drug relapse, seeking was reinstated for 20 minutes by exposure to light and tone cues previously paired with cocaine delivery, but no cocaine was delivered. Both olanzapine- and risperidone-treated animals pressed the active lever significantly more than vehicle-treated animals over the course of the reinstatement session, again indicating increased magnitude of craving and impaired extinction learning (**Fig. 1D-E**).

**Figure 1.**
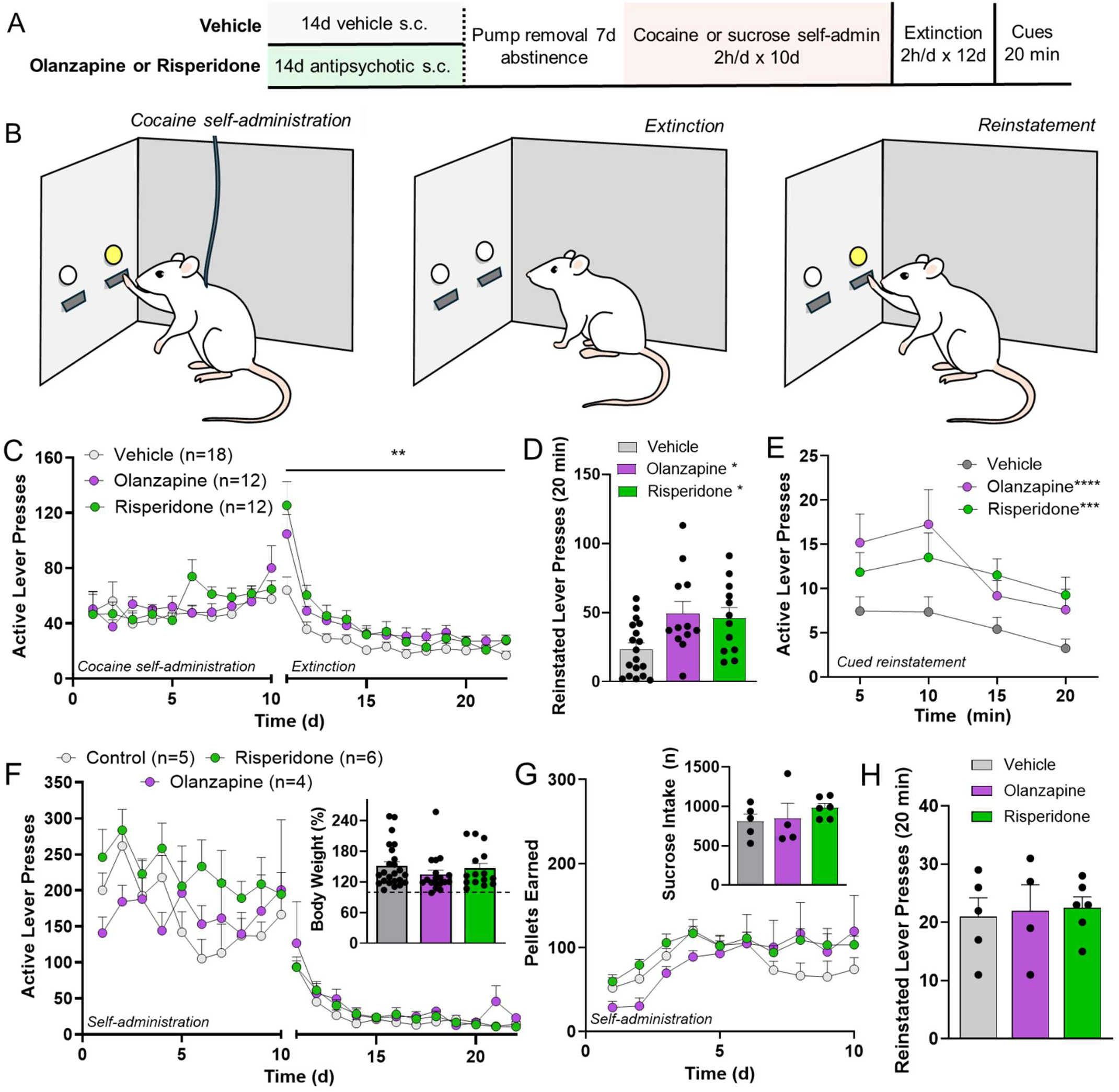
Chronic olanzapine or risperidone treatment increases cocaine-seeking in rat. (**A**) Adult Long Evans rats received subcutaneous infusion of olanzapine (2.5 mg/kg/d), risperidone (4 mg/kg/d) or vehicle for 14 days via osmotic infusion. After pump removal and a 7-day washout period, rats were trained to self-administer i.v. cocaine or oral sucrose pellets during daily sessions, and reward delivery was paired with light and tone cues (**B**). Animals then underwent extinction training, where reward and cue delivery were withheld in response to active lever pressing (**B**, extinction). (**C**) Lever pressing was then reinstated by exposure to light and tone cues for 20 minutes. (**C**) Antipsychotic-treated rats exhibited delayed extinction learning relative to vehicle-treated animals and continued to press the active lever for cocaine despite reward absence. During cued reinstatement, antipsychotic-treated rats also pressed significantly more for cocaine (**D**) and continued pressing for longer (**E**) despite no reward delivery. Instead, antipsychotic-treated rats trained to self-administer sucrose did not differ in active lever pressing during self-administration or extinction (**F**), body weight (**F**, inset), sucrose intake (**G**) or reinstated sucrose seeking (**H**) compared with vehicle-treated control rats.

### Chronic Treatment with Second-Generation Antipsychotics Did Not Impact Natural Reward Intake or Seeking

To determine whether antipsychotic pretreatment increased intake of or craving for non-pathological rewards, the same experimental design was repeated using oral sucrose pellets as a reinforcer. Animals pretreated with antipsychotics did not differ in body weight gain in the time between osmotic pump implantation and self-administration (**Fig. 1F, inset**) or in active lever pressing during sucrose self-administration or extinction (**Fig. 1F**), suggesting that antipsychotic pretreatment did not increase craving or impair extinction of contextual or cue-based associations involving natural rewards. Over the course of self-administration, rats did not differ in the amount of sucrose obtained (**Fig. 1G**) and active lever pressing during a 20-min cue-induced reinstatement session was also not different between treatment groups (**Fig. 1H**), suggesting that the previously observed impairment in reversal learning was selectively induced by the addictive reinforcer cocaine in antipsychotic-treated animals.

### Elevated Relapse Risk after Antipsychotic Discontinuation Was Driven by D2-MSN Plasticity in the Ventral Striatum

Elevated reward-seeking during cue-induced reinstatement has been linked previously with post-synaptic potentiation of D1 receptor-expressing medium spiny neurons (D1-MSNs) in the nucleus accumbens core (NAcore), part of the ventral striatum, measured as increased spine head diameter due to AMPA-type glutamate receptor insertion (41, 42). Instead D2-MSNs have been found to undergo potentiation during extinction learning, where reward seeking is gradually reduced through removal of a reinforcer (42). To determine whether the elevated reward-seeking in antipsychotic-treated rats was driven by these well-established canonical mechanisms, we virally-labeled D1- or D2-MSNs in the NAcore by delivery of AAV2-CAG-Flex-Ruby2sm-Flag in D1- or D2-Cre rats (**Fig. 2A**). Individual dendritic segments from D1- or D2-MSNs were imaged and quantified after 20 minutes of cue-induced reinstatement of cocaine seeking in vehicle- or antipsychotic-treated animals (**Fig. 2B**). We found that D1-MSNs in the NAcore of animals treated with either olanzapine or risperidone exhibited smaller dendritic spines relative to vehicle-treated animals (**Fig. 2C**), suggesting that D1-MSN potentiation did not contribute to elevated cocaine seeking in these animals. Instead, spine head diameter was significantly increased in NAcore D2-MSNs in animals pretreated with olanzapine or risperidone (**Fig. 2D**), suggesting antipsychotic treatment elicited neuronal potentiation selectively in this population of neurons. No differences in D1- or D2-MSN spine density were observed between treatment groups (**Fig. S2**).To determine whether the potentiation of D2-MSNs contributed to elevated reward seeking after antipsychotic treatment, we delivered a designer receptor exclusively activated by designer drugs (DREADD) to NAcore D2-MSNs using AAV2-hSyn-DIO-hM4D(Gi)-mCherry in D2-Cre rats before training them to self-administer cocaine. After extinction training, lever pressing was reinstated by exposure to drug-associated cues and NAcore D2-MSNs were inhibited by i.p. delivery of the designer drug clozapine N-oxide (CNO). Antipsychotic-treated rats that received an inert intracranial infusion (saline) and CNO prior to cued reinstatement exhibited elevated lever pressing compared to vehicle-treated, antipsychotic-naïve rats (**Fig. 2E**), as previously observed (**Fig. 1D-E**). D2-MSN inhibition in DREADD-expressing animals that received CNO reversed the increase in lever pressing conferred by antipsychotic pretreatment (**Fig. 2E**), confirming that D2-MSN potentiation in the NAcore contributed to elevated cocaine-seeking in antipsychotic-treated rats.

**Figure 2.**
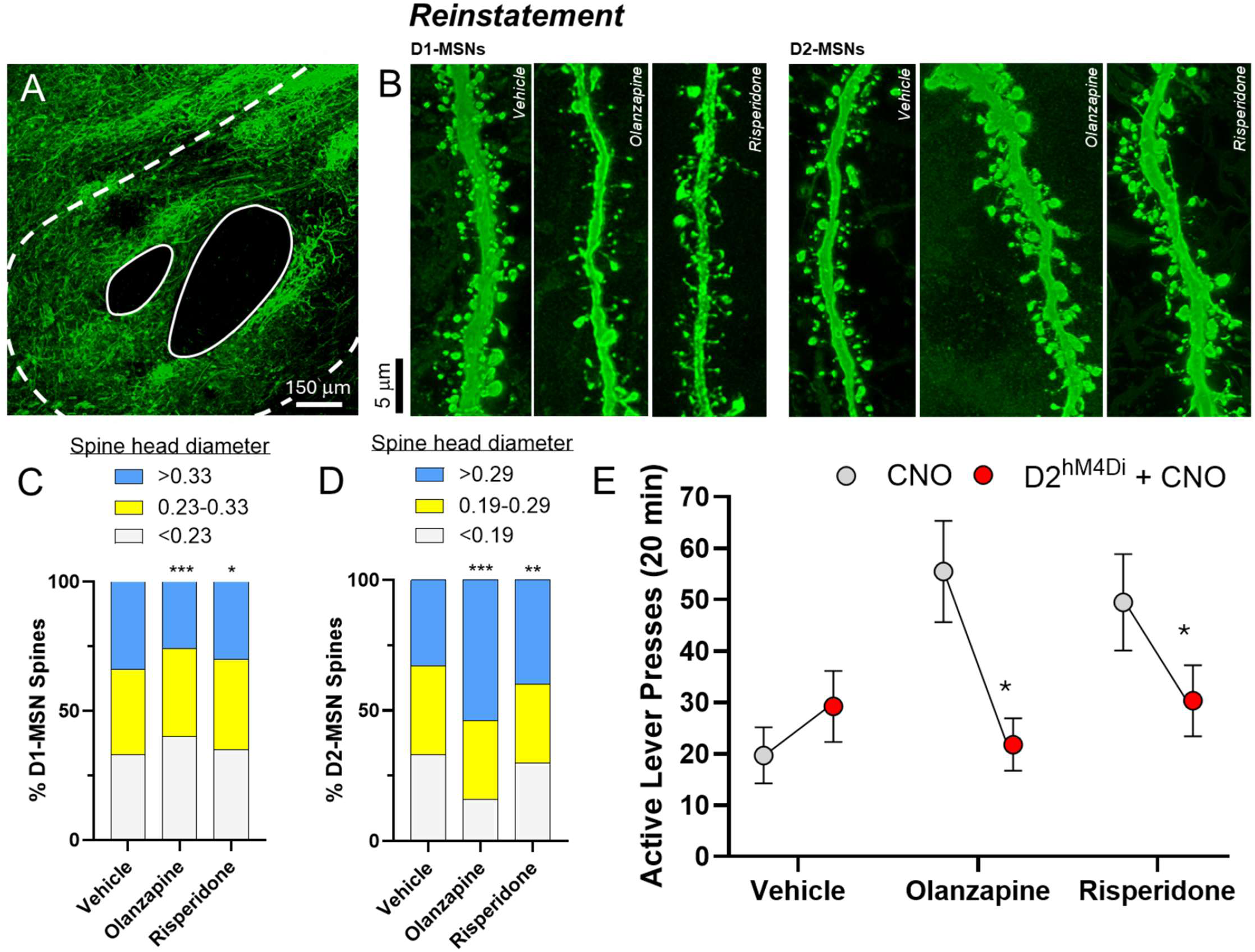
Antipsychotic-induced cocaine relapse vulnerability was driven by NAcore D2-MSN plasticity. (**A**) AAV2-CAG-Flex-Ruby2sm-Flag was delivered to the NAcore of D1- and D2-Cre rats and diameter of dendritic spines on D1- and D2-MSNs was quantified during 20 minutes of reinstated cocaine seeking in antipsychotic- or vehicle-treated rats (**B**). Olanzapine and risperidone treatment resulted in a relative decrease in spine head diameter on D1-MSNs (**C**) and a corresponding increase in spine head diameter on D2-MSNs during reinstated cocaine seeking (**D**). (**E**) AAV2-hSyn-DIO-hM4D(Gi)-mCherry was delivered to the NAcore in D2-Cre rats and viral incubation occurred over the course of operant training. Saline was infused intracranially instead of AAV2-hSyn-DIO-hM4D(Gi)-mCherry in control animals (**E**, gray circles) and CNO was delivered prior to cued reinstatement of cocaine seeking in all rats. DREADD inhibition of D2-MSNs (**E**, red circles) reversed the antipsychotic-induced increase in active lever pressing during cued reinstatement.

### Extinction Training Did Not Elicit D2-MSNs Plasticity in Antipsychotic-Treated Rats

To determine whether D2-MSN potentiation was present in antipsychotic pretreated animals prior to relapse, we repeated D1- and D2-MSN labeling and quantitation during 20 minutes of extinction training 24 hours after the last cocaine self-administration session (**Fig. 3A**). While antipsychotic treatment did not impact spine density on D1- or D2-MSNs during extinction training, treatment with either olanzapine or risperidone reduced spine head diameter on D1-MSNs compared to vehicle treatment (**Fig. 3C**), similar to what was observed during cued reinstatement testing. Olanzapine, but not risperidone treatment resulted in an overall decrease in D2-MSN spine head diameter during extinction training relative to vehicle (**Fig. 3D**).

**Figure 3.**
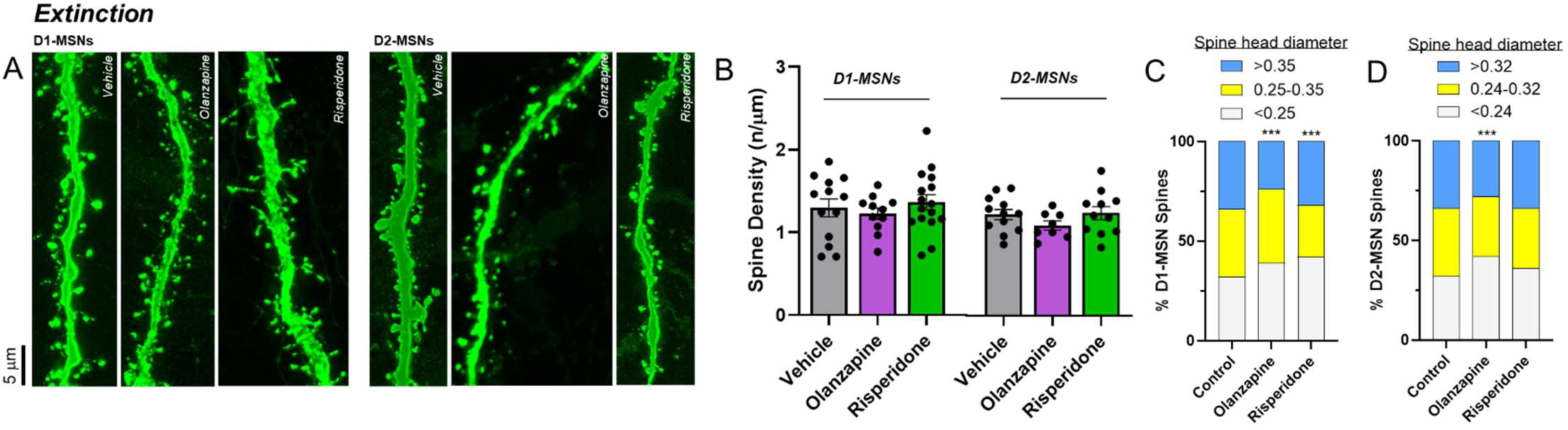
D2-MSN spine head diameter increased selectively during cued cocaine seeking. (**A**) AAV2-CAG-Flex-Ruby2sm-Flag was delivered to the NAcore of D1- and D2-Cre rats and spine density and head diameter were quantified during 20-minutes of extinction training 24h after the last self-administration session. **(B)** Antipsychotic treatment did not impact spine density on D1- or D2-MSNs. Olanzapine and risperidone treatment decreased spine head diameter on D1-MSNs during extinction training (**C**). Only olanzapine decreased D2-MSN spine head diameter during extinction training.

### Duration of Antipsychotic Treatment Predicted SUD Severity

To determine whether these observations were consistent with findings observed in humans, and to determine whether the antipsychotic or addictive drug type, or treatment dosing or duration impacted the pattern we observed in our rodent model, we performed a systematic review of all literature where antipsychotic medications were used to treat addictive substance use or craving. We screened 7907 studies collected from CENTRAL, Embase, Scopus and PubMed and found 61 studies that met our inclusion criteria (**Fig. S3**). Of these, 20 studies reported no change in addictive substance craving or use following antipsychotic treatment, 14 reported an increase in craving or use after antipsychotic treatment compared to placebo, and 27 reported a decrease in craving or use after antipsychotic treatment compared to placebo. Antipsychotic dose ranges were similar across studies and many of the same antipsychotic drugs were used in studies that reported increases and decreases in addictive substance craving or use (**Fig. 4A**). Category of addictive substance used (i.e. alcohol, psychostimulant, opioid, or cannabis) did not impact whether a study reported increased or decreased addictive substance craving or use. However, antipsychotic treatment duration strongly predicted study outcome and studies using shorter antipsychotic treatment timelines were more likely to report decreased addictive substance craving or use (**Fig. 4C**). For example, more than half of studies where antipsychotic treatment reduced drug craving or use delivered antipsychotics for 2 weeks or fewer. Instead, nearly half of studies reporting an increase in craving or use after antipsychotic treatment used treatment timelines of 4 months or more. Treatment timecourses between 1-3 months were equally represented across categories, and were the most predominant timecourse used among studies that reported no change in addictive substance craving or use (**Fig 4C**). Because aripiprazole relies on a unique mechanism compared to other antipsychotic drugs, including olanzapine and risperidone used in our animal studies, partially agonizing, rather than antagonizing the D2r, we repeated the same analysis after excluding studies of aripiprazole, and found the same reliance of study outcome on antipsychotic treatment duration (**Fig. S4**).

**Figure 4.**
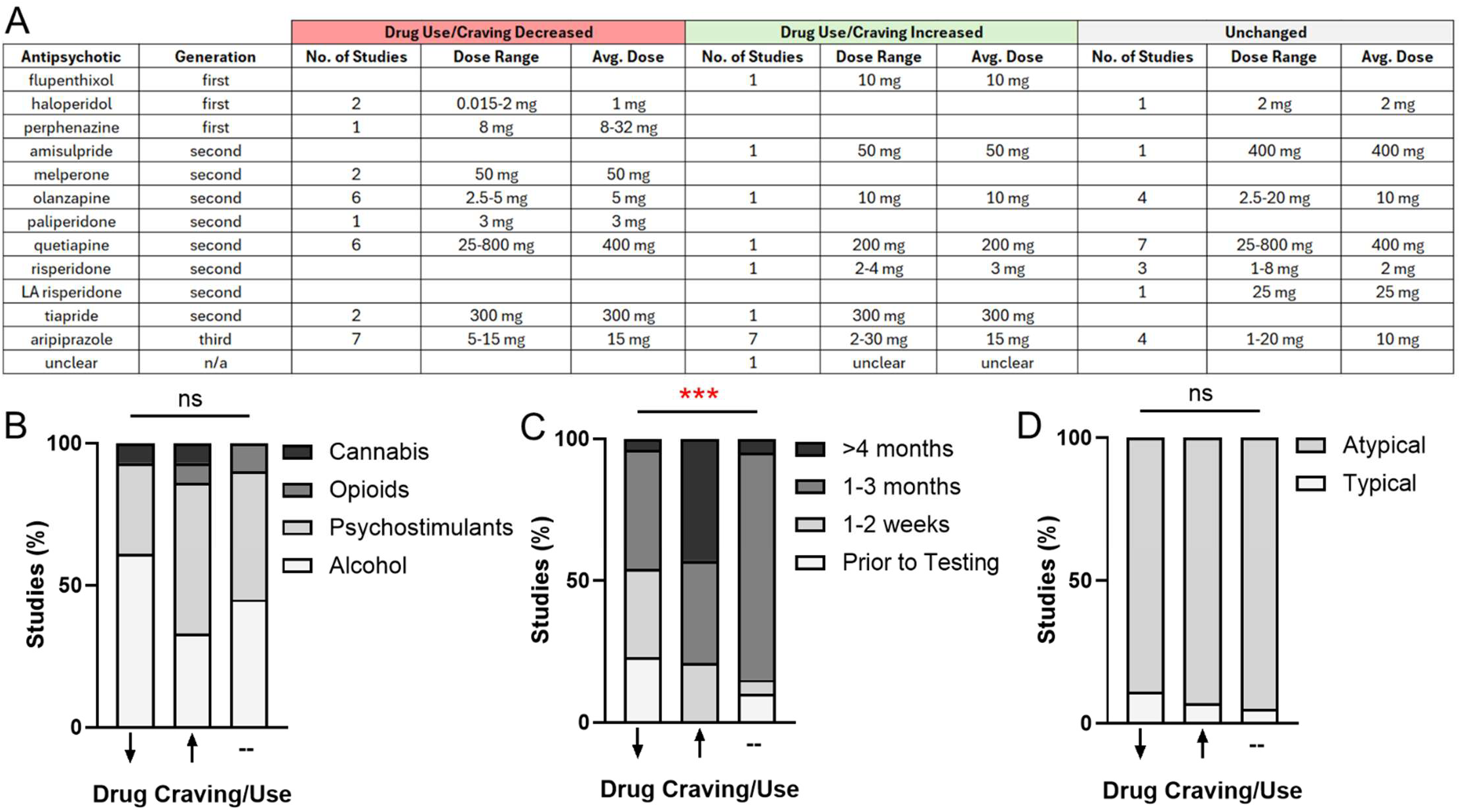
Chronic antipsychotic treatment lasting 4 months of more increased addiction vulnerability in patients. (**A**) Clinical studies examining the impact of antipsychotic treatment on measures of addiction vulnerability were analyzed by systematic review. Similar doses of antipsychotic medications were given across studies and did not impact whether studies found increased, decreased, or unchanged measures of addictive substance craving or use. (**B**) The type of misused substance (alcohol, psychostimulants, opioids or cannabis) also did not affect whether or how antipsychotic treatment impacted addiction measures. (**C**) Instead, duration of antipsychotic treatment strongly impacted whether a study reported an increase or decrease in addictive substance craving or use, with treatment timelines of 4 months or longer over-represented in studies reporting an increase in addiction vulnerability measures. (**D**) The antipsychotic used (first, second or third generation) also had no impact on whether studies reported increases or decreases in addictive substance craving or use.

## Discussion

Antipsychotics are approved for schizophrenia, bipolar disorder, depression, autism, and Tourette syndrome, and are prescribed off-label for insomnia, agitation, hyperactivity, and anxiety in adolescents and adults (8, 43–48). While often effective, both first- and second-generation antipsychotics can be accompanied by significant side effects when given chronically, with first-generation antipsychotics triggering motor symptoms including dyskinesias and second-generation medications resulting in metabolic changes that lead to weight gain and obesity (49). We recently characterized the complex neurobiology underlying some of these side effects following chronic treatment with, and discontinuation of, the first-generation antipsychotic haloperidol and identified increased cocaine relapse in a rat model of intravenous cocaine self-administration as an additional adverse effect (30). The current study was designed to assess whether SUDs vulnerability also results from treatment with second-generation antipsychotics and how these findings compare to data from patients. Collectively both animal and human findings reported here extend our previous observations (30) and demonstrate that chronic treatment with antipsychotics across drug classes, including first- and second-generation antipsychotics, as well as the third-generation antipsychotic aripiprazole, confer similar vulnerability to SUDs, whether or not treatment was discontinued as in our previous and current animal studies.

### Animal Study

Using a rat model of cocaine use disorder, we found that two weeks of treatment with second-generation antipsychotics, followed by a brief discontinuation period, impaired extinction of cocaine seeking during withdrawal and enhanced cue-induced cocaine relapse, both well-established behavioral markers of addiction vulnerability (50–52). The increase in motivated behavior in antipsychotic-treated rats was selective for addictive, but not natural rewards, since antipsychotic-treated animals did not display reversal learning deficits during extinction of operant sucrose self-administration or elevated sucrose seeking during reinstatement. Mechanistically, our data indicate that chronic antipsychotic treatment elevates drug seeking through potentiation of D2-MSNs in the NAcore, extending our previous findings that first-generation antipsychotics drive locomotor sensitization to cocaine through D2-MSN synaptic potentiation (30). This is notable because addiction-related behaviors are more commonly linked to D1-MSN potentiation, whereas D2-MSNs are often associated with extinction learning and aversion (42, 53). Antipsychotic-treated rats also exhibited impaired extinction learning, reflected by persistently elevated lever pressing during extinction of cocaine self-administration, yet this deficit was not observed after sucrose self-administration. Thus, the extinction impairment observed in antipsychotic-treated rats appears selective for drug rewards. Interestingly, although dynamic D2-MSN activity has been linked to reversal learning (54, 55), we found no evidence of D2-MSN plasticity during 20 minutes of extinction training in antipsychotic-treated rats relative to controls. Moreover, while olanzapine reduced D2-MSN spine head diameter during extinction, risperidone impaired extinction without affecting spine morphology, suggesting that additional mechanisms contribute to these extinction deficits. How prior cocaine exposure recruits D2-MSN potentiation after antipsychotic treatment, and how this potentiation selectively promotes drug seeking without broadly disrupting reversal learning, remain open questions. One possibility is that antipsychotic pretreatment potentiates a specific subpopulation of D2-MSNs linked to motivation, as suggested by some prior work, including our own (30).

Importantly, both male and female rats that self-administered cocaine showed extinction deficits after antipsychotic treatment, although females exhibited a more robust increase in cued reinstatement (**Fig. S1**). These findings may be relevant in light of evidence that several adverse effects of antipsychotics, including weight gain, hyperprolactinemia, and cardiac alterations, are often more severe in female patients (56). We also note that chronic antipsychotic treatment in rodents does not appear to alter D2 receptor expression in the NAcore (30), arguing against receptor upregulation as the mechanism underlying D2-MSN potentiation and increased drug seeking. Because D2 receptors are inhibitory, compensatory upregulation would be expected to suppress, rather than potentiate, D2-MSN activity. The receptor mechanisms responsible for the reduced dopaminergic signaling (37) and enhanced glutamatergic signaling (30) observed during chronic antipsychotic treatment therefore remain to be identified and represent an important direction for future work.

An important question raised by these findings is how 14 days of antipsychotic exposure used in rodents relates to clinically relevant treatment timelines in humans. Because antipsychotic clearance is substantially faster in rodents than in patients, continuous drug delivery over 14 days likely models a much longer period of exposure in humans (57, 58). Continuous delivery by osmotic minipump was therefore used to better approximate clinically relevant pharmacokinetics and sustained receptor occupancy (57, 58). On this basis, the 14-day treatment window used here may approximate 6–7 months or more of treatment in humans (59, 60), a duration that aligns with the treatment range associated with poorer SUD outcomes in the clinical literature according to our systematic review and meta-analysis (**Fig. 4**). Consistent with previous work, prolonged antipsychotic exposure in rodents produced striatal synaptic adaptations associated with addiction vulnerability (29), whereas shorter exposures appear less likely to do so and may even suppress drug seeking (61, 62).

In the present study, olanzapine and risperidone were administered at 2.5 mg/kg/day and 4 mg/kg/day, respectively, doses that fall within the clinically relevant range once interspecies differences in body weight and metabolism are taken into account. Given their differing affinities for the D2 receptor (approximately 59.3 nM for olanzapine and 4.6 nM for risperidone), the risperidone regimen likely produced relatively high D2 receptor blockade, whereas olanzapine produced lower D2 receptor blockade. Notably, antipsychotics that produce increased addiction vulnerability in our rodent model to date, including haloperidol, olanzapine, and risperidone, share high affinity for dopamine D2–D4 receptors, modest affinity for D1 receptors, and high affinity for selected adrenergic and serotonergic receptors, although the precise receptor activity that recruits D2-MSN plasticity and drives drug seeking is unknown.

### Human Study

The clinical literature showed that the strongest predictor of addiction vulnerability across studies was the duration of antipsychotic treatment. Studies using acute or very short treatment schedules (single exposure or daily treatment for 2 weeks or fewer) were more likely to report improvements in craving or substance use outcomes. Intermediate treatment durations (1–3 months) did not consistently favor either beneficial or detrimental outcomes. In contrast, antipsychotic treatment lasting 4 months or longer was significantly more likely to be associated with increased drug craving, more frequent positive urine screens, and/or greater addictive substance use, regardless of antipsychotic class or addictive substance type. These findings align closely with the translational interpretation of our rodent data, in which 14 days of continuous antipsychotic exposure may approximate a prolonged treatment window in humans as discussed above.

By contrast, antipsychotic dose did not emerge as a consistent predictor of SUD severity across clinical studies. Doses reported in the literature ranged from 2.5–20 mg/day for olanzapine and 1–8 mg/day for risperidone (**Fig. 4)**, suggesting that treatment duration, rather than dose alone, may be the more critical determinant of addiction-related outcomes. Although many of the clinical studies included both male and female participants, sex was often not balanced within treatment groups and data were rarely reported in a way that allowed assessment of whether variables such as antipsychotic dose or duration differentially affected addiction vulnerability in men and women. Similarly, prior antipsychotic exposure could not be controlled for in patient populations requiring chronic treatment before study entry. For this reason, patients with preexisting psychiatric illness treated chronically with antipsychotics were not included here. Nonetheless, given the high rates of comorbidity between psychiatric disorders and SUDs (2–4), it is plausible that the relationship between chronic antipsychotic treatment and addiction vulnerability may extend across diagnostic groups.

Taken together with our previous work, these findings suggest that both first- and second-generation antipsychotics may increase addiction vulnerability when administered over extended periods. This may also apply to non-canonical antipsychotics such as aripiprazole, which has likewise been associated with increased alcohol or drug use and craving after prolonged treatment (**Fig. 4**). Importantly, the present findings should not be interpreted as arguing against antipsychotic use, which remains essential for many patients. Rather, they highlight the need to better understand how individual antipsychotic drugs influence reward-related plasticity so that treatment strategies can be optimized to minimize unintended effects on motivation and addictive substance use while preserving therapeutic benefit. This issue is particularly relevant given the broad and increasing use of antipsychotics not only for schizophrenia, but also for mood and anxiety disorders, agitation, and insomnia (63–65), including in adolescents (43, 44), a population that may be especially vulnerable to long-lasting neuroplastic effects of pharmacological intervention.

In conclusion, our animal and human findings indicate that chronic antipsychotic treatment can increase vulnerability to addictive substance use across drug classes. Our systematic review and meta-analysis identified treatment duration as the key variable associated with addictive substance use and relapse outcomes. In rodents, this effect is mediated by potentiation of D2-MSNs in the NAcore, highlighting a cellular substrate for antipsychotic-associated addiction liability. Importantly, the effect does not appear to be explained by D2 receptor blockade alone, but instead by downstream neuroadaptations induced by prolonged treatment. Clinical data further suggest that this vulnerability emerges not only after discontinuation, but also during ongoing treatment, particularly when exposure lasts 4 months or longer. Testing new molecules with antipsychotic effects, including novel muscarinics and non-dopaminergic agents (66, 67), for their impact on SUDs remains a priority along with refinement of available treatments to avoid or counteract the observed effects on SUDs severity in those requiring chronic treatment.

## Acknowledgements

This work was supported by the Brain & Behavior Research Foundation (NARSAD Young Investigator Award to D.A.), the University of Cincinnati (Innovation Award to D.A.) and the National Institutes of Health (DA054339 to A.K.).

## Supplementary Figures and Legends

**Figure S1.**
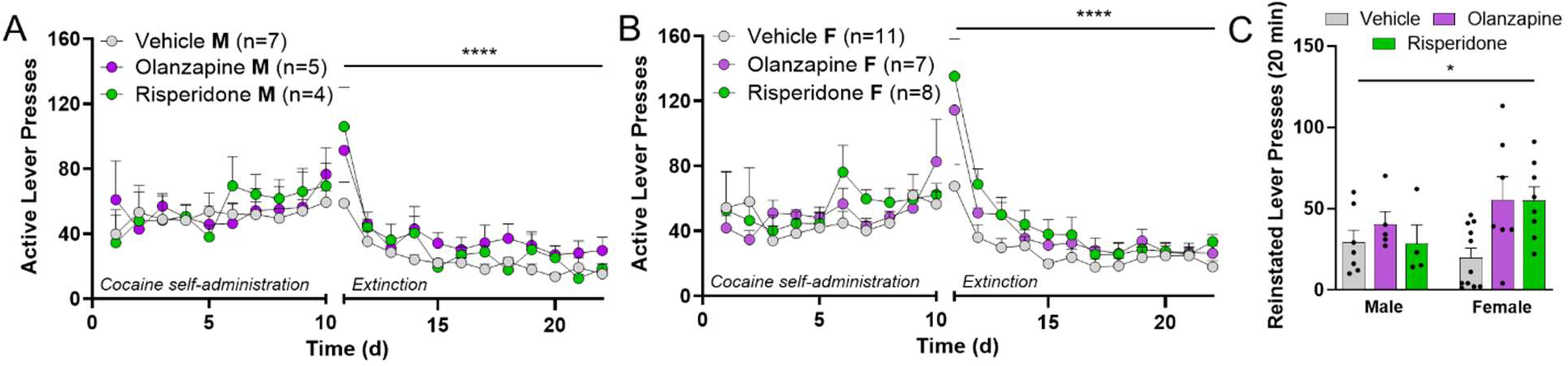
Antipsychotic-induced extinction deficits were observed in male and female rats. Both male (**A**) and female (**B**) rats exhibited extinction deficits after treatment with second-generation antipsychotics olanzapine or risperidone. (**C**) Elevated lever pressing during cue-induced reinstatement of cocaine seeking was more pronounced in female rats. (**A**-**B**) ****p<0.0001 vs. Vehicle by 2-way ANOVA, (**C**) *p<0.05 effect of treatment by 1-way ANOVA.

**Figure S2.**
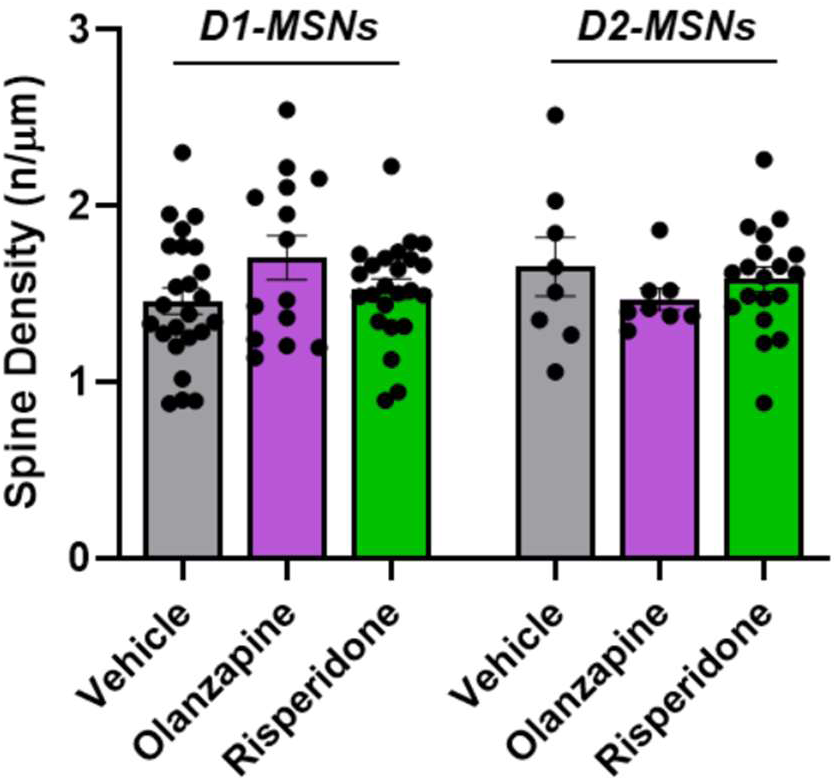
Spine density during cued reinstatement was unchanged by antipsychotic pretreatment. Spine density on D1- and D2-MSNs was unchanged by pretreatment with either olanzapine or risperidone.

**Figure S3.**
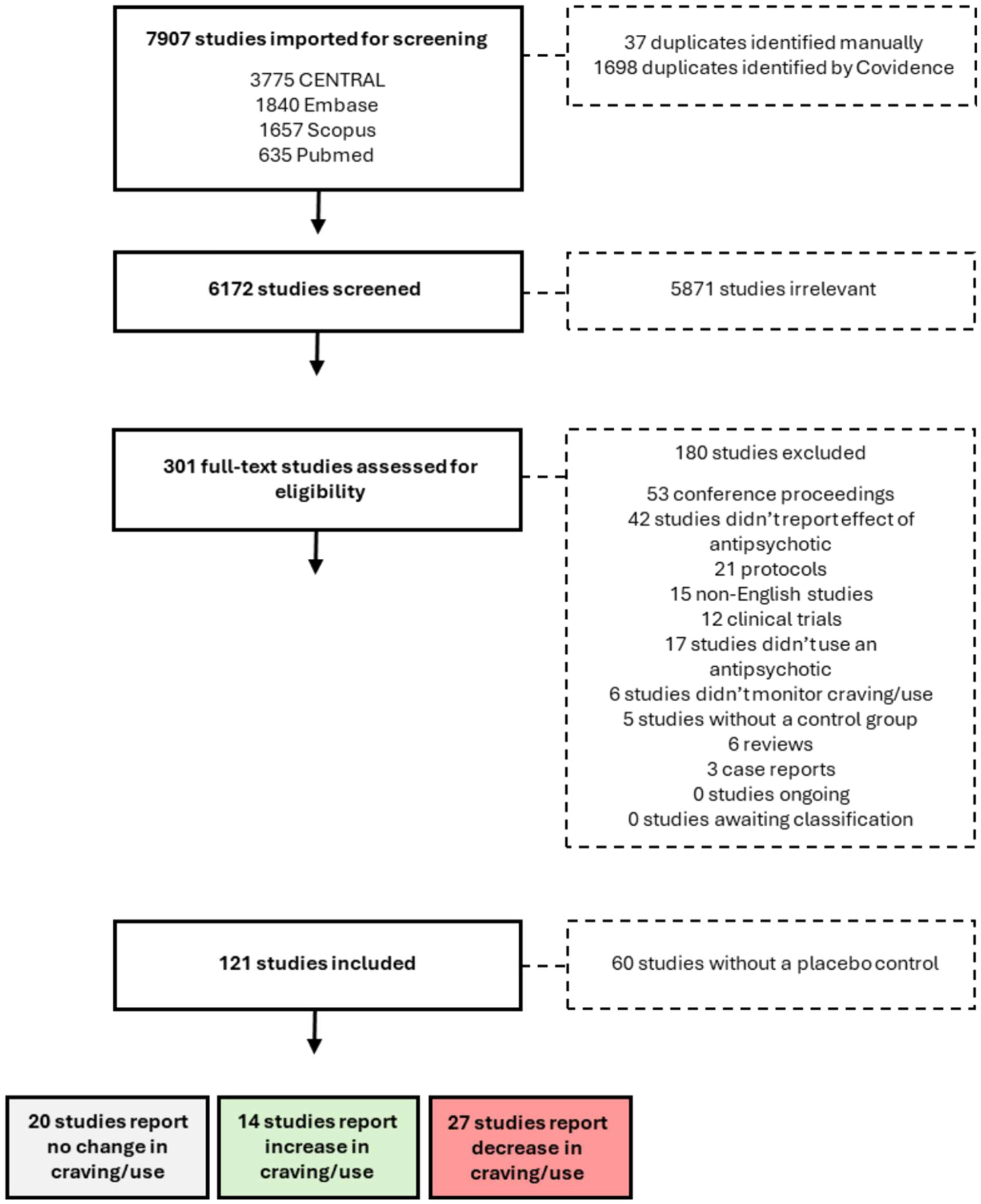
PRISMA flow diagram of systematic review. A total of 7907 papers were imported from CENTRAL, Embase, Scopus and PubMed. After removal of duplicates and addition of manually-identified articles, 6172 papers underwent title and abstract screening. Of these, ultimately 301 papers were selected for full text review. After removal of irrelevant publications and studies lacking placebo controls, a total of 61 papers were ultimately analyzed for effects of antipsychotic treatment on addictive substance craving and/or use.

**Figure S4.**
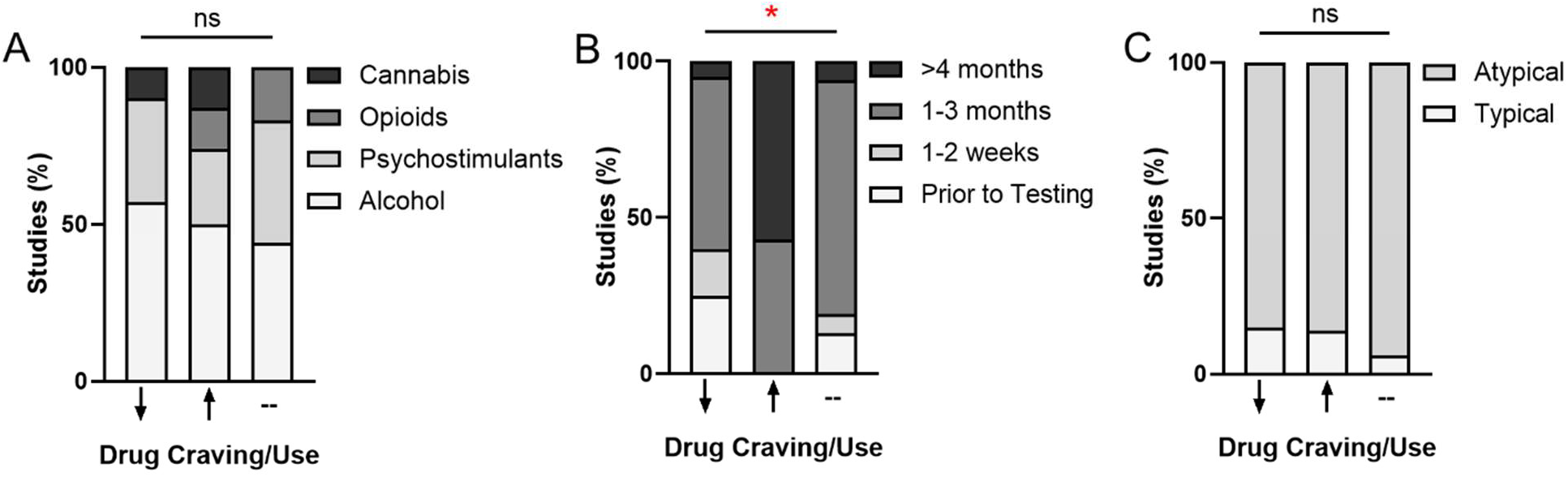
Chronic antipsychotic treatment increased addictive substance craving and/or use in patients. (**A**) Type of substance use disorder (i.e. cannabis, opioid, stimulant or alcohol) did not impact whether publications reported decreased (downward arrow), increased (upward arrow), or unchanged (dash) addictive substance craving or use following antipsychotic treatment. (**B**) Treatment duration significantly impacted whether studies reported increased, decreased or unchanged addictive substance craving or use, with antipsychotic treatment lasting 4 months or longer increasing addiction severity in patients (*p<0.05 using Chi^2^). **(C)** Type of antipsychotic used for treatment (typical, first generation vs. atypical, second or third generation) did not impact study outcomes.

